# Hippocampal and caudate network specificity is altered in older adults

**DOI:** 10.1101/2022.07.14.500121

**Authors:** Michael V. Freedberg

## Abstract

The hippocampal and the caudate networks, defined by their intrinsic resting state functional connectivity (FC), exhibit strong network specificity. This is reflected as strong within-network and weak between-network FC, as well as their dissociable roles in cognition. Aging typically reduces the specificity of brain networks. However, whether the hippocampal and caudate networks show age-related decreases in network specificity is unclear. Further, whether any age-related decreases in network specificity are due to reduced within-network FC, increased between-network FC, or some combination of both, is also unclear. Using a large-scale fMRI data set acquired from healthy younger (*n* = 101, aged 18–40 years) and older (*n* = 101, 55–75 years) adults, networks centered on the left anterior hippocampus and the head of the right caudate nucleus were compared. These sub-regions were chosen based on their distinct contributions to cognition, their known interactions, and their susceptibility to age-related changes. A mixed effect model was used to identify brain regions where network specificity differed between groups. For younger adults, hippocampal network specificity was strong in the medial frontal gyrus (MFG), while caudate network specificity was strong in the basal nuclei. However, network specificity in these same regions was significantly reduced in older adults. Reduced specificity in each network was due to a weakening of within-network connectivity rather than an increase in between-network connectivity. These results indicate that hippocampal and caudate network specificity decreases with advancing age, raising the possibility that these reductions may contribute to age-related changes in memory.

**Impact Statement:** Older adults represent an increasing proportion of the population, thus increasing the urgency for us to understand the neurological changes that contribute to age-related memory loss. Aging is associated with a loss of network specificity, reflected as reduced within-network, and increased between-network, connectivity. My work shows that this phenomenon is not restricted to the hippocampal and caudate networks, which are critical for learning facts and skills, respectively: Both networks decreased in network functional connectivity. These results raise the possibility that “blurring” between these networks relates to age-related memory loss, possibly as a compensatory mechanism to protect memory in old age.

## 1. INTRODUCTION

The hippocampus and the caudate and their intrinsically connected networks play mostly dissociable roles in memory. For example, episodic memory (i.e., memory for personally-experienced and verbally recallable information; Tulving, 1987) depends heavily on neural activity within a hippocampus-centered brain network that includes the medial prefrontal cortex, posterior cingulate, precuneus, and angular gyri (Ranganath and Ritchey, 2012; Ritchey et al., 2015). In contrast, procedural memory (i.e., memory for skills and habits; Keele et al., 2003; Yin and Knowlton, 2006) is closely tied to activity within a caudate-centered brain network that includes the ventrolateral thalamus and the dorsolateral prefrontal cortex (Alexander and Crutcher, 1990; Haber and Calzavara, 2009; Hikosaka et al., 1995; Knowlton et al., 1996; Poldrack et al., 1999). Although several key studies have reported that activity within each of these networks is largely independent (Knowlton et al., 1996; Saint-Cyr et al., 1988; Scoville and Milner, 1957), other work shows that these networks instead inhibit one another (Freedberg et al., 2022; Poldrack et al., 2001; Wimmer et al., 2014). That is, activation of the hippocampus-centered network suppresses activation of the caudate-centered network, and vice versa. Further, the extent to which these two brain networks interact is associated with memory performance (Brown et al., 2012; Dickerson et al., 2011; Freedberg et al., 2022; Mattfeld and Stark, 2010; Moses et al., 2010).

In cognitively healthy younger adults without memory impairments, the hippocampal and caudate networks show high network specificity. This is reflected as strong within-network functional connectivity (FC) and weak between-network FC (Achard et al., 2006; Achard and Bullmore, 2007; Stam, 2004). The hippocampal network exhibits strong resting state FC (rsFC) with the bilateral angular gyri, medial parietal cortex, and frontal cortex (Fox et al., 2005; Fox and Raichle, 2007), while the caudate network exhibits strong rsFC with the superior, middle and frontal gyri, basal nuclei, anterior cingulate cortex, posterior cingulate cortex, inferior/medial temporal gyri, and inferior parietal lobule (Di Martino et al., 2008). These findings strongly suggest that high network specificity is a hallmark of the healthy young adult brain. In contrast, network specificity in older adults is reduced (Andrews-Hanna et al., 2007; Chan et al., 2014; Damoiseaux et al., 2008; Geerligs et al., 2015; Malagurski et al., 2020; Oschmann et al., 2020; Spreng et al., 2016; Staffaroni et al., 2018; Yang et al., 2014; Zhang et al., 2021). In the default mode (Cao et al., 2014; Staffaroni et al., 2018; Yang et al., 2014), sensorimotor (Malagurski et al., 2020), frontoparietal (Geerligs et al., 2015; Oschmann et al., 2020), salience (Oschmann et al., 2020), and dorsal attention (Spreng et al., 2016) networks, this reduced network specificity is due to reduced within-network connectivity (Andrews-Hanna et al., 2007; Chan et al., 2014; Damoiseaux et al., 2008; Geerligs et al., 2015; Oschmann et al., 2020; Spreng et al., 2016; Staffaroni et al., 2018; Yang et al., 2014; Zhang et al., 2021), increased between-network connectivity (Chan et al., 2014; Geerligs et al., 2015; Malagurski et al., 2020; Spreng et al., 2016; Zhang et al., 2021), or a combination of both (Chan et al., 2014; Geerligs et al., 2015; Spreng et al., 2016; Zhang et al., 2021).

Previous work has reported that the mutual inhibition between the hippocampal and caudate networks is reduced in older adults, such that these two networks cooperate to support memory (Tomás Pereira et al., 2015; Voss et al., 2018). These findings suggest that the strong hippocampal-caudate network specificity observed in healthy younger adults diminishes with increasing age. To date, patterns of hippocampal and caudate rsFC have never been directly compared between younger and older adults. Therefore, aging-related changes in hippocampal-caudate network specificity are unknown. Here, I address this gap in knowledge by directly comparing hippocampal and caudate rsFC patterns between younger and older adults. I reason that if aging decreases network specificity of the hippocampal and caudate networks, then the extent to which the hippocampus and caudate are functionally coupled with their respective network nodes should be lower in older than in younger adults. For example, let us assume that a given brain region (e.g., the medial prefrontal cortex; MFG) is more strongly functionally coupled to the hippocampus than to the caudate in younger adults. An age-related decline in the network specificity of this brain region could be due to decreased coupling of this region with the hippocampus (reduced within-network connectivity; Figure 1, left), increased coupling of this region with the caudate (increased between-network connectivity; Figure 1, middle), or some combination of the two (Figure 1, right).

**Figure 1.**
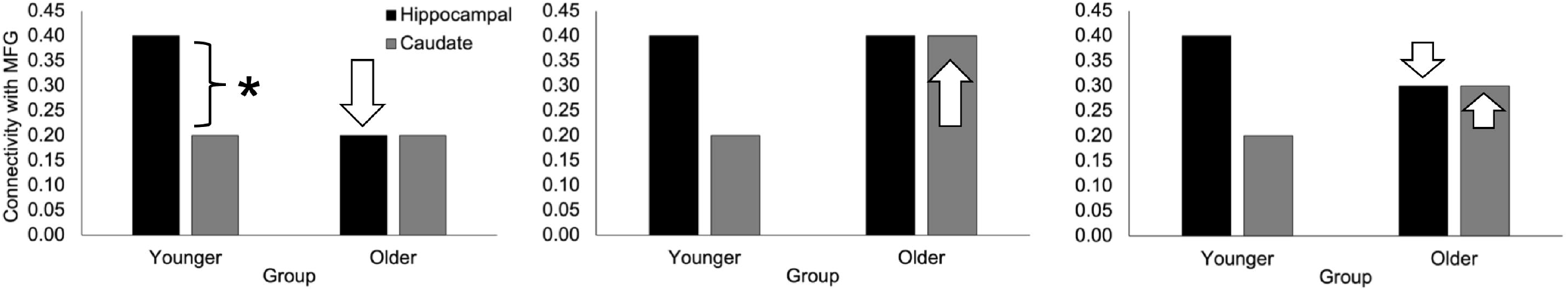
Hypothetical sources of reduced hippocampal network specificity in older adults. Network specificity is represented as the difference in connectivity between the hippocampus and the medial frontal gyrus (MFG; asterisk). Loss of network specificity can occur due to a loss of within-network functional coupling (left), an increase in between-network functional coupling (middle), or a combination of the two (right).

The primary objective of the current work was to determine whether hippocampal and caudate network specificity is altered in older adults, and if so, to characterize these alterations. Given that hippocampal-mediated memory (i.e., episodic memory) exhibits stronger age-related declines than caudate-mediated memory (i.e., procedural memory; Mitchell et al., 1990; Vakil and Agmon-Ashkenazi, 1997), I predicted that age-related decline in network specificity would be apparent within the hippocampal network, but not the caudate network.

## 2. METHODS

### 2.1. Overview

Imaging data were obtained from the Enhanced Nathan Kline Institute’s Rockland Sample (NKI-RS; http://fcon_1000.projects.nitrc.org/index.html). The NKI-RS is a large-scale online repository that includes neuroimaging data from a diverse age group (7–85 years) of community-dwelling New Yorkers (Nooner et al., 2012). Anonymized functional neuroimaging data from the NKI-RS are publicly available for download. Only structural and resting-state scans were used in the present study. The study was approved by the NKI Institutional Review Board, and all participants provided their informed consent.

### 2.2. Participants

From the entire NKI-RS sample (*N* > 1,000), 242 younger (aged 18–40 years) and 169 older (55–75 years) right-handed adults were identified. Owing to data storage constraints, and to better equate sample sizes, the younger adult sample was reduced by 124 by including only odd-numbered younger adult participants. Fifty-four older and 17 younger adults were excluded because they did not have a useable functional or anatomical data file (old = 8, young = 7), they had excessive head motion during the resting-state scan (old = 45, young = 8), and/or parcellation of their anatomical scan failed (old = 1, young = 2). This left 101 younger and 115 older adults. These groups differed significantly in numbers of male and female participants (*X*^2^ = 5.86, *p* < 0.05). To balance groups by number and sex (*X*^2^ = 3.01, *ns*), 14 random older females were removed from the sample. Thus, the final sample included 101 younger adults (mean age = 27.18 ± 5.83; 50 females) and 101 older adults (mean age = 63.84 ± 5.28; 73 females). Figure 2 provides the flowchart for participant exclusion/inclusion.

**Figure 2.**
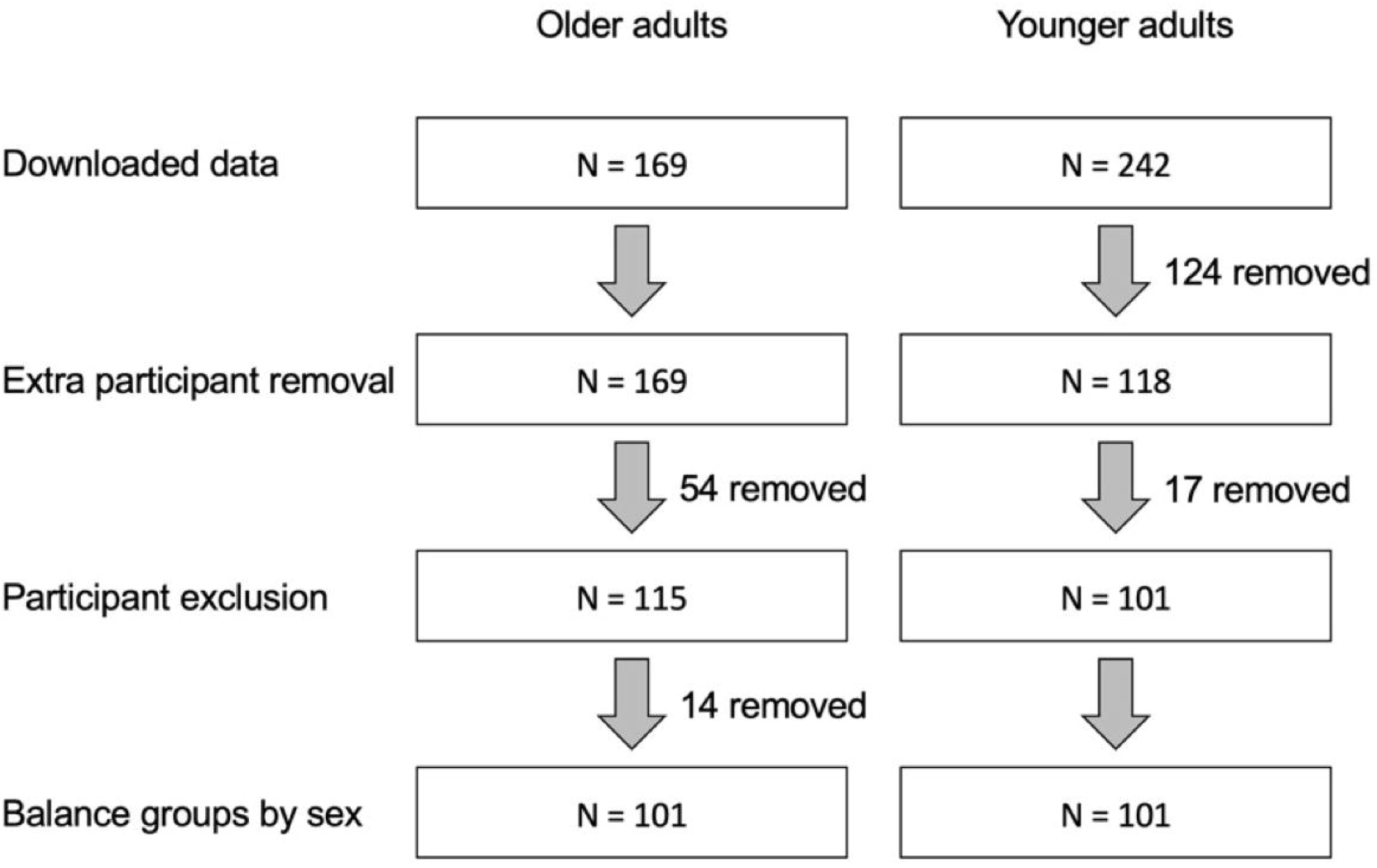
Flowchart of participant exclusion/inclusion.

### 2.3. MRI data acquisition and processing

Resting-state scans with three different scanning parameters are available as part of the NKI-RS data set: (1) TR = 645 ms, voxel size = 3 mm^3^, scan length = ~10 min; (2) TR = 1,400 ms, voxel size = 2 mm^3^, scan length = ~10 min, (3) TR = 2,500 ms, voxel size = 3 mm^3^, scan length = ~5 min. The second set was used because of the smaller voxel size, which maximizes spatial resolution, and because longer scans produce more reliable FC estimates (Birn et al., 2013). Resting-state data were acquired using a Siemens Magnetom Trio 3T MRI scanner while participants kept their eyes open and stared at a fixation cross. Blood oxygen level-dependent (BOLD) data were recorded using an echoplanar imaging sequence (EPI: TR = 1,400 ms, TE = 30 ms, flip angle = 65°, 64 transversal contiguous interleaved slices per volume, 2.0 slice thickness, FOV 2.24 cm^2^, voxel size = 2 mm^3^, volumes = 404; scan length 9.43 min). A magnetization-prepared rapid gradient echo sequence (MPRAGE; TR = 1,900 ms, TE = 2.50 ms, 176 slices per volume, 1 mm thickness, FOV = 25.6 cm^2^, 256 × 256 acquisition matrix, voxel size = 1.0 mm isotropic, 4.30 min) was used for structural/functional co-localization during image processing and to identify seed regions.

Resting-state data were preprocessed with Analysis of Functional Images software (AFNI version 21.1.03; Cox, 1996; RRID:SCR_005927). Preprocessing included removal of the first five volumes (*3dTcat*), de-spiking (*3dDespike*), slice-timing correction to the first slice (*3dTshift*), deobliquing (*3dWarp*), motion correction (*3dvolreg*), functional/structural affine coregistration to Talairach space using the TT_N27 anatomical template (*align_epi_anat.py* and *3dAllineate*), spatial smoothing using a 4 mm full width at half maximum (FWHM) Gaussian kernel (*3dmerge*), scaling the voxel time series to a mean of 100, with a range of 0–200 (*3dTcat* and *3dcalc*), head motion regression from each voxel time series using the mean and derivatives of six parameter estimates (pitch, roll, yaw, and rotation around each axis), and high-pass filtering via linear detrending (*3dDeconvolve*). The estimates for motion correction, EPI to anatomical alignment, and transformation into standard space were combined into a single transformation so that regridding of EPI data occurred only once. Head motion correction, filtering, and linear detrending were performed based on a single regression model in order to avoid reintroducing signal related to nuisance covariates, caused by projecting data into a sequence of different subspaces (Lindquist et al., 2019). Frames that included movement displacement greater than 0.3 mm were censored prior to model regression (*3dDeconvolve, 3dTproject, 3dcalc*). As mentioned above, 45 older and 8 younger adult participants were excluded for exceeding an average head displacement of 0.3 mm across all frames.

### 2.4. rsFC calculations

The left anterior hippocampus and the head of the right caudate are hubs in the episodic and procedural memory networks, respectively. The encoding of recent episodic memories is specifically associated with activation of the left anterior hippocampus (Moscovitch et al., 2006; Zeidman and Maguire, 2016), and neuromodulation targeted to the left anterior hippocampus enhances episodic memory (Freedberg et al., 2022; Hermiller et al., 2020; Kim et al., 2018; Nilakantan et al., 2017; Wang et al., 2014). In contrast, the head of the right caudate is activated during nonmotor (Poldrack et al., 1999) and motor (Doyon et al., 1996) procedural learning. Additionally, these areas are commonly observed to interact with each other (Freedberg et al., 2020), suggesting that age-related changes in hippocampal and caudate network rsFC may be interdependent. Thus, I defined the hippocampal and caudate networks based on rsFC centered on these structures due to their distinct functional roles.

A mask of the left anterior hippocampus was constructed by parcellating each participant’s anatomical scan with the “recon-all” command in FreeSurfer version 7.1.0 (Dale et al., 1999) and the Desikan atlas (Desikan et al., 2006). To constrain each participant’s hippocampal mask to the anterior aspect, all left hippocampal voxels posterior to Talairach y-coordinate y = 19 were discarded. This coordinate was used because it corresponds to the uncal apex on the CA_N27_ML atlas left hippocampus in TT_N27 template space. The uncal apex is commonly used as the major anatomical point dividing the anterior and posterior hippocampus (Brunec et al., 2018; Poppenk et al., 2013). A mask of the head of the right anterior caudate was also constructed for each participant using FreeSurfer. The right caudate nucleus (CA_N27_ML atlas) was limited to the anterior one third of caudate voxels in each participant. Finally, this region of interest was distinguished from the ventral striatum by discarding voxels with z-direction values less than seven (Di Martino et al., 2008; Postuma and Dagher, 2006). rsFC centered on each region of interest was calculated by averaging the time series of all voxels within each mask and then regressing that time series against the time series of all brain voxels. All *R*-values were Fisher *Z*-transformed to normalize hippocampal and caudate rsFC.

### 2.6. Statistical analyses

Hippocampal and caudate rsFC maps were submitted to a 2 x 2 linear mixed-effects model. The results were limited to a group mask that excluded white matter and ventricles. The effects of group (young and old) and network (hippocampal and caudate) and their interaction were modeled (*3dLMEr*). A random effect of participant was included to isolate participant-specific variability. Average head displacement per frame [young: 0.16 ± 0.07 mm; old: 0.18 ± 0.06 mm; *t*(200) = 3.99, *p* < 0.001] and temporal signal to noise ratio [young: 73.37 ± 15.30; old: 82.13 ± 17.74; *t*(200) = 3.76, *p* < 0.001] differed significantly between groups. Although motion regression and other preprocessing steps help reduce the influence of such differences on FC analyses (Power et al., 2014) these values were included as covariates to reduce any residual influence on the results. Because these covariates differed between groups, covariates were centered based on group averages (Chen, 2014; Chen et al., 2014). All analyses were thresholded at *p* < 0.001 and corrected with a false discovery rate (FDR) of *q* < 0.05. Post hoc analyses were performed on surviving clusters. The results of the main effects are shown using results from *3dttest++*, which outputs positive and negative *t*-values. These analyses did not use covariates. Multiple comparisons correction among clusters was performed by controlling the FDR using Benjamini and Hochberg’s method (Benjamini and Hochberg, 1995).

To determine whether these results were driven by differences in the sex of participants between groups, a control analysis was performed. For each participant, caudate FC maps were subtracted from hippocampal FC maps and fed into a 2 x 2 linear mixed-effects model using group (young vs. old) and sex as factors. The results were thresholded using the same approach described above (*p* < 0.001, *q* < 0.05, cluster size = 40).

## 3. RESULTS

Whole brain rsFC values centered on the left anterior hippocampus and the head of the right caudate nucleus were submitted to a group (old vs. young) and network (hippocampal vs. caudate) linear mixed effects model. It was expected that the hippocampal but not the caudate network would show reduced network specificity in older adults compared to younger adults. In my analysis, this would be represented by a significant interaction between group and network as well as post hoc analyses showing a reduced difference in hippocampal vs. caudate connectivity within the hippocampal network. Although the critical part of the analysis is the interaction between group and network, the main effects of each are described below for completeness.

### 3.1. Age-related effects on rsFC (main effect of group)

Table 1 shows regions where rsFC differed between age groups irrespective of network (hippocampal and caudate). This analysis revealed 11 clusters where young adults showed significantly greater rsFC with the hippocampal and caudate memory network hubs (i.e., the anterior hippocampus and head of the right caudate, respectively) than older adults. These included subcortical and somatomotor regions, such as the left and right caudate, postcentral gyri, and the cerebellum. All regions shown in Table 1 survived multiple comparisons correction.

**Table 1.** Clusters showing significant group (age) effects (young > old; *p* < 0.001, *q* < 0.05, cluster size = 40). Columns 2–4 show the coordinates of the voxel demonstrating the peak difference between groups. Columns 5–6 show the region where the cluster showed the highest overlap with any region in the CA_N27_ML atlas and the percent overlap value. Effect size is shown in the last column.

| Voxels | Peak x | Peak y | Peak z | Region of Maximal Overlap | % Overlap | Cohen's *d* |
| --- | --- | --- | --- | --- | --- | --- |
| 402 | 7 | 13 | 4 | Right caudate nucleus | 54.1 | 0.96 |
| 394 | 21 | -25 | 62 | Right postcentral gyrus | 43.5 | 0.67 |
| 345 | -3 | 7 | 2 | Left caudate nucleus | 80.3 | 0.85 |
| 339 | -29 | -23 | 62 | Left postcentral gyrus | 42.3 | 0.69 |
| 165 | 45 | 9 | -2 | Right insula lobe | 62.3 | 0.85 |
| 131 | 45 | -43 | -24 | Right cerebellum (Crus 1) | 29.0 | 0.82 |
| 108 | -21 | -5 | 2 | Left putamen | 41.5 | 0.69 |
| 85 | -41 | 17 | 0 | Left inferior frontal gyrus | 61.4 | 0.85 |
| 84 | -17 | 9 | -6 | Left amygdala | 32.7 | 0.80 |
| 61 | -1 | -19 | 2 | Left thalamus | 10.8 | 0.77 |
| 42 | 33 | -3 | 56 | Right middle frontal gyrus | 87.7 | 0.60 |

### 3.2. Regional differences in rsFC within age groups (main effect of network)

The main effect of network contrasted hippocampal and caudate rsFC irrespective of group. This aspect of the analysis reveals regions that are more tightly associated with the hippocampal network (i.e., regions that show a bias toward hippocampal rsFC) and regions that are more tightly associated with the caudate network (i.e., regions that show a bias toward caudate rsFC). At a significance level of *p* < 0.001 (FDR corrected, cluster size = 40), most voxels within the group mask showed significance. However, increasing the p-value to 0.0001 revealed remarkable correspondence in the pattern of rsFC bias between the two age groups (Figure 3).

**Figure 3.**
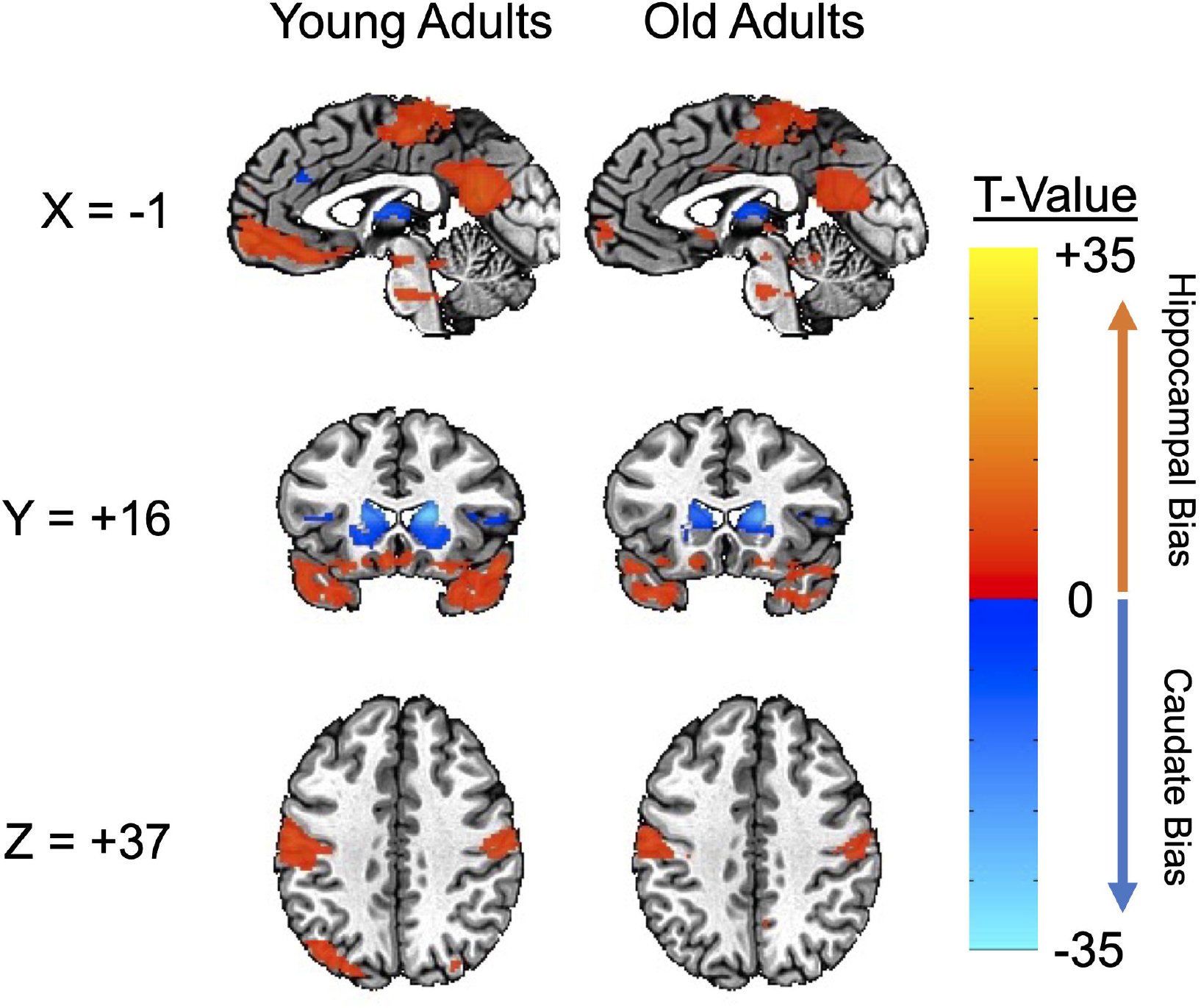
Clusters showing significantly different hippocampal vs. caudate rsFC for younger and older adults (3dttest++, *p* < 0.0001, *q* < 0.05, cluster size = 40, no covariates). Warm colors indicate stronger hippocampal rsFC; cool colors indicate stronger caudate rsFC.

### 3.3. Age-related differences in resting-state hippocampal and caudate rsFC (group by network interaction)

The interaction between group and network revealed regions where network specificity differed between younger and older adults. Network specificity was defined as a reduced difference in the strength of hippocampal and caudate connectivity within their respective networks. Table 2 shows clusters of significance for the interaction between group and network. Of the five clusters, four indicated reduced network specificity in the caudate network; these were mostly localized to basal nuclei regions (bilateral caudate and putamen). The final cluster showed reduced network specificity in the hippocampal network and was localized to the left MFG (see Figure 4).

**Figure 4.**
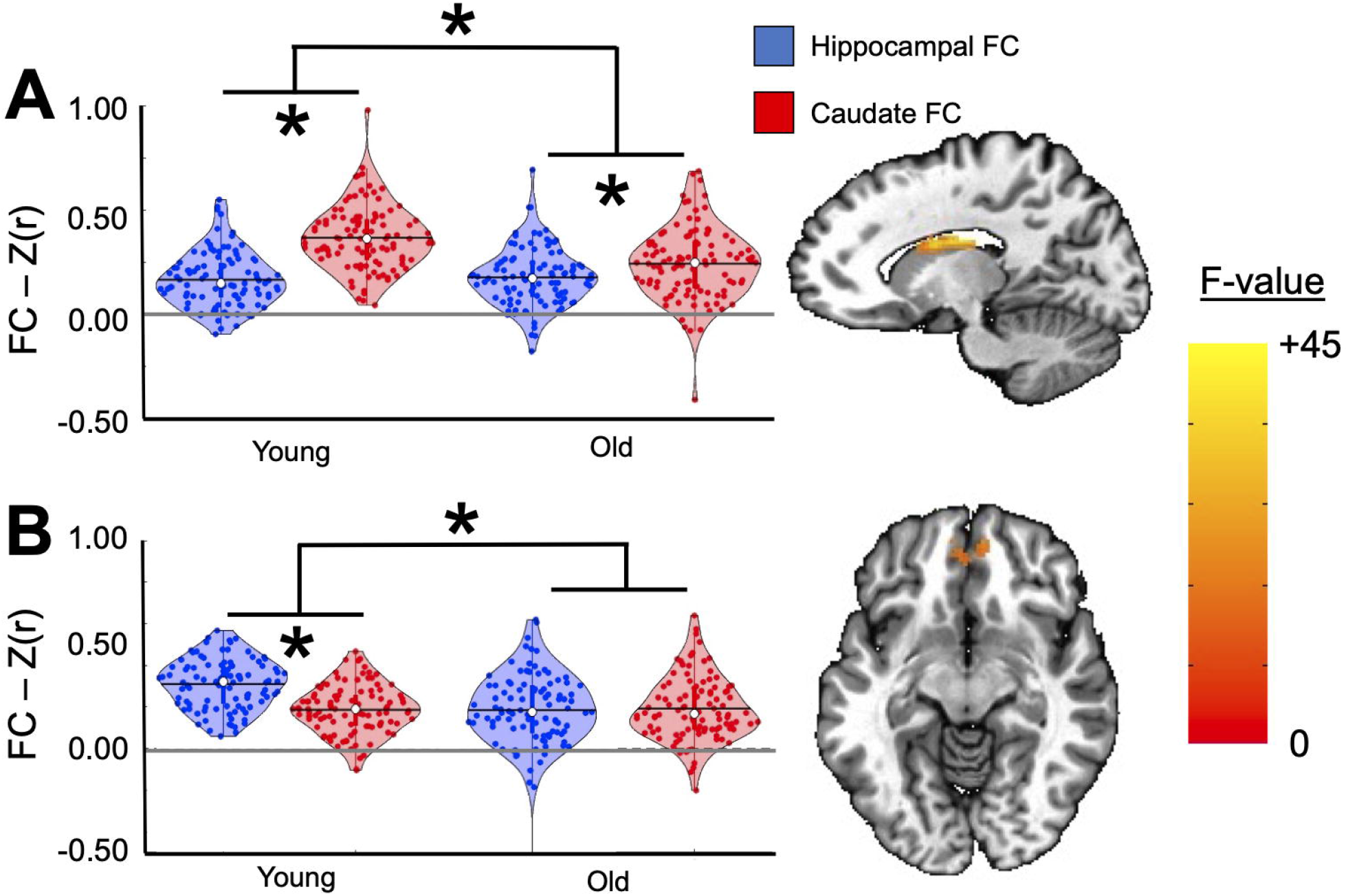
Violin plots showing significant interaction between group and network. Left hippocampal (blue) and right caudate (red) rsFC with the left caudate (A) and medial frontal gyrus (B; *3dLMER, p* < 0.001, *q* < 0.05, cluster size = 40) are shown. Dots represent individual participants. Black lines represent means; white dots represent median. Grey lines represent zero. * *p* < 0.001.

**Table 2.** Significant clusters from group by network interaction (*3dLMER, p* < 0.001, *q* < 0.05, cluster size = 40). Columns 2–4 show the coordinates of the voxel demonstrating the peak interaction estimate. Columns 5–6 show the region where the cluster showed the highest overlap with any region in the CA_N27_ML atlas and the percent overlap value. Effect size is shown in the last column. Clusters 1-4 show evidence of reduced network specificity in the caudate network, while Cluster 5 shows reduced specificity in the hippocampal network.

| Cluster | Voxels | Peak x | Peak y | Peak z | Region of Maximal Overlap | % Overlap | Cohen's $d$ |
| --- | --- | --- | --- | --- | --- | --- | --- |
| 1 | 402 | 13 | -9 | 22 | Right caudate nucleus | 79.7 | 0.86 |
| 2 | 345 | -13 | -9 | 22 | Left caudate nucleus | 80.4 | 0.85 |
| 3 | 339 | -17 | 11 | -4 | Left putamen | 55.9 | 0.93 |
| 4 | 165 | 17 | 9 | -4 | Right putamen | 56.2 | 0.68 |
| 5 | 394 | 7 | 49 | -6 | Left medial frontal gyrus | 32.7 | 0.87 |

Separate post hoc *t*-tests were performed contrasting hippocampal and caudate rsFC for each group. These tests were used to determine whether network specificity was observable at the five significant clusters for each group. Younger adults displayed strong network specificity. The difference between hippocampal and caudate rsFC was highly significant for all clusters [right caudate nucleus: *t*(100) = 16.00, *p* < 0.001, *d* = 1.24; left medial frontal gyrus: *t*(100) = 9.00, *p* < 0.001, *d* = 0.93; left caudate nucleus: *t*(100) = 14.91, *p* < 0.001, *d* = 1.16; left putamen: *t*(100) = 11.19, *p* < 0.001, *d* = 0.88; right putamen: *t*(100) = 9.97, *p* < 0.001, *d* = 0.79]. For older adults, network specificity was only observable at two of the clusters, including the right [*t*(100) = 6.36, *p* < 0.001, *d* = 0.63] and left caudate nucleus [*t*(100) = 3.45, *p* < 0.001, *d* = 0.40]. Network specificity was not observable at the left medial frontal gyrus [*t*(100) = 0.15, *p* = .88, *d* = 0.02], left putamen [*t*(100) = 1.35, *p* = .18, *d* = 0.15], and right putamen [*t*(100) = 0.12, *p* = .90, *d* = 0.01]. All significant post hoc contrasts survived multiple comparisons correction. The control analysis examining the effect of sex revealed no evidence that sex interacted with group, nor was any main effect of sex observed.

## 4. DISCUSSION

The left anterior hippocampus (Knowlton et al., 1996; McCormick et al., 2015; Zeidman and Maguire, 2016) and head of the right dorsal caudate (Poldrack et al., 1999; Saint-Cyr et al., 1988) are critical hubs in the episodic and procedural memory networks, respectively. Based on their distinct roles in cognition and their high network specificity in younger adults, I asked whether network specificity is reduced in the hippocampal and caudate networks (as defined by their intrinsic patterns of resting functional connectivity) in older versus younger adults. Although I predicted that the hippocampal, but not the caudate, network would show a loss of network specificity, the results show a loss of network specificity in both networks: younger adults showed significantly stronger hippocampal network connectivity with a cluster in the MFG and significantly stronger caudate network connectivity with bilateral regions of the basal nuclei, whereas these biases were significantly reduced in older adults. Furthermore, hippocampal network connectivity with a cluster in the medial frontal gyrus was so reduced in older adults that it was statistically indistinguishable from caudate connectivity. Likewise, caudate network connectivity with the left and right putamen was so reduced in older adults that it was statistically indistinguishable from hippocampal connectivity. Thus, despite previous literature showing that procedural memory is largely preserved in older adults (Clark et al., 2015; Vakil and Agmon-Ashkenazi, 1997), the caudate network showed significant evidence of reduced network specificity. Finally, in both networks, reduced network specificity was due to a drop in the strength of connectivity within networks rather than an increase in connectivity with the other network (Figure 4).

The aging brain undergoes numerous structural and functional changes that could explain the loss of network specificity observed here. One candidate is the natural loss of dopamine neurons (5–10% every decade) that occurs with aging (Fearnley and Lees, 1991). Dopamine midbrain neurons project to both networks (Chowdhury et al., 2013; Gasbarri et al., 1994; Graybiel and Ragsdale, 1979) and are theorized to mediate interactions between networks (Freedberg et al., 2020; Goto and Grace, 2005b). This is supported by several neuroimaging studies in younger adults (Braskie et al., 2011; Nyberg et al., 2016; Wimmer et al., 2014; Wimmer and Shohamy, 2012; Wittmann et al., 2005). For example, dopamine-related learning signals in the striatum are related to the consolidation of episodic memories in the hippocampus several weeks later (Wittmann et al., 2005). Additionally, reward prediction errors, which evoke dopamine neurotransmission (Schultz et al., 1997), interfere with episodic memory formation (Wimmer et al., 2014). Finally, in older adults, the density of striatal dopamine transporters predicts episodic memory decline (Rieckmann et al., 2018). These studies suggests that age-related loss of dopamine neurons may contribute to dysregulation of hippocampal and caudate network activity.

Alternatively, age-related changes in the structure (Salat et al., 1999) and function (Grady, 2008) of the prefrontal cortex may be associated with decline in hippocampal and caudate network specificity (Freedberg et al., 2020). The prefrontal cortex interacts with the striatum and the hippocampus during learning (Antzoulatos and Miller, 2014, 2011; Goto and Grace, 2005b, 2005a; Puig et al., 2014). For example, direct stimulation of the prefrontal cortex directly causes antagonistic interactions between the striatum and the hippocampus (Goto and Grace, 2005a). If the prefrontal cortex becomes less efficient with age, then this antagonistic influence on striatal-hippocampal interactions may cause more cooperative interactions between these networks, possibly contributing to reduced network specificity.

Although the key finding from the present study is that network specificity is decreased in older adults, general group-level differences in FC were also observed. In general, rsFC was significantly greater in younger than in older adults. These differences occurred in regions within both networks, including the MFG. Interestingly, when the results were split by network, the head of the left caudate showed a significant decrease in both hippocampal and caudate rsFC in older adults compared with younger adults. This suggests that this region may be especially vulnerable to aging. Finally, the pattern of network connectivity was remarkably similar between groups (Figure 3). However, this aspect of the analysis was qualified by the significant interaction, which indicated a significant loss of network specificity for older adults.

The large sample size afforded by the NKI-RS online repository (Nooner et al., 2012) is a strength of the present study. The most significant limitation is the reliance on rsFC. The results do not indicate how network specificity is altered in older adults when networks are engaged during task performance. For example, Voss et al. (2018) found that older adults engage both hippocampal and striatal networks during memory performance while young adults only engage the hippocampal one (Voss et al., 2018). These results, however, were not observed when examining patterns of rsFC. Thus, it is possible that network specificity is especially reduced, or even possibly maintained, during task performance.

## 5. CONCLUSION

In this study, I asked whether aging is associated with reduced network specificity within the hippocampal and caudate networks. Hippocampal and caudate rsFC patterns obtained from a large dataset were contrasted between younger and older adults. Contrary to expectations, both the hippocampal and caudate networks showed evidence of reduced network specificity. Medial frontal gyrus connectivity was highly specific to the hippocampal network in younger adults but was statistically indistinguishable from the caudate network in older adults. Likewise, basal nuclei connectivity was highly specific to the caudate network in younger adults but was significantly reduced in older adults. Loss of hippocampal and caudate network specificity was due to significantly reduced within-network connectivity, but not increased between-network connectivity. These results indicate that aging is associated with reduced hippocampal and caudate network specificity, raising the possibility that these reductions may contribute to age-related changes in memory.

## Acknowledgements

The author acknowledges the Texas Advanced Computing Center (TACC) at The University of Texas at Austin for providing HPC resources that have contributed to the research results reported within this paper. URL: http://www.tacc.utexas.edu.

## Conflict of Interest Statement

I declare no conflict of interest.

## Funding Statement

This research did not receive any specific grant from funding agencies in the public, commercial, or not-for-profit sectors.

## Notes

### Competing Interest Statement

The authors have declared no competing interest.

